# Dynamics of anteroposterior axis establishment in a mammalian embryo-like system

**DOI:** 10.1101/2021.02.24.432766

**Authors:** Kerim Anlaş, Nicola Gritti, David Oriola, Krisztina Arató, Fumio Nakaki, Jia Le Lim, James Sharpe, Vikas Trivedi

**Affiliations:** EMBL Barcelona, 08003 Barcelona, Spain; Institució Catalana de Recerca i Estudis Avançats, 08010, Barcelona, Spain; EMBL Heidelberg, Developmental Biology Unit, 69117 Heidelberg, Germany

## Abstract

In the mammalian embryo, specification of the anteroposterior (AP) axis demarcates one of the first steps of body plan formation. While this process requires interactions with extra-embryonic tissues in the native embryo, minimal *in vitro* systems from embryonic stem cells (ESCs) undergo initial AP polarization in the absence of any localized, external cues. This self-organizing potential of stem cells remains not well understood. Here, we study such an initial symmetry breaking event in gastruloids, an established *in vitro* model for mammalian body plan formation, using the mesodermal marker gene Brachyury or T (Bra/T) to denote the onset of AP axis specification and concomitant germ layer formation. Through aggregate fusion experiments and manipulation of initial culture conditions as well as key developmental signalling pathways, we probe the dynamics of Bra/T polarization. We further conduct single-cell (sc) RNA sequencing of gastruloids at early stages to identify incipient molecular signatures of germ layer commitment and differences between Bra/T^+^ and Bra/T^−^ populations during as well as after symmetry breaking. Moreover, we transcriptionally compare early development of gastruloids to the mouse embryo and conclude that gastruloids reproducibly undergo AP axis and germ layer specification in a parallel, but distinct manner: While their primed pluripotent cell populations adopt a more mesenchymal state in lieu of an epithelial epiblast-like transcriptome, the emerging mesendodermal lineages *in vitro* are nevertheless similar to their in vivo equivalents. Altogether, this study provides a comprehensive analysis of self-organized body plan establishment in a minimal *in vitro* system of early mammalian patterning and highlights the regulative capacity of mESCs, thereby shedding light on underlying principles of axial polarity formation.

## 2. Introduction

The bilaterian body plan is established during early embryogenesis as part of a process commonly referred to as gastrulation, whereby collective cell rearrangements give rise to a multilayered blueprint with three germ layers (ecto-, meso- and endoderm) organized along three body axes: anteroposterior (AP), dorso-ventral (DV) and mediolateral (ML) [1, 2, 3].

These events have been extensively studied in several organisms across the animal kingdom and the mouse embryo is a prominent example in which the AP axis is the first to be newly determined, whereas DV axial polarity is technically inherited from the Blastocyst’s embryonic-abembryonic axis [4, 5]. AP axis specification is initiated by a subpopulation of the extraembryonic primitive endoderm (PrE), the distal visceral endoderm (DVE) precursor cells, at embryonic day (E)4.5. At E5.5, the DVE proper undergoes migration to the prospective anterior region, giving rise to the anterior visceral endoderm (AVE), expressing markers including Wnt and Nodal inhibitors. The primitive streak is formed by E6.5 at the diametrically opposite end, thereby demarcating the posterior, characterized by activity of Wnts, Nodal and Bra/T [6, 7, 8].

Although particularly mammalian early body plan patterning is thus tightly coupled to biochemical and mechanical clues from extraembryonic tissues, findings in (embryo-like) *in vitro* systems illustrate that AP axis formation can occur in an autonomous manner [9, 10, 11, 12].

In this context, gastruloids from aggregated mouse embryonic stem cells (mESCs) represent a prime example: They are grown without any localized signalling inputs, yet nevertheless develop comparatively advanced axial organization, congruent with the early mouse embryo [13, 14]. After a few days in non-adherent culture, such aggregates undergo elongation and display spatially delimited expression of marker genes for the AP, DV and ML axes, concomitant with the emergence of distinct cell populations analogous to the three germ layers [15].

This apparent self-organizing capability of ESCs cannot be observed *in vivo* due to the presence of extra-embryonic tissues and species-specific stereotypical development, but may provide clues about underlying principles of mammalian body plan formation which cannot be revealed by studying only the native embryo [16, 17]. Investigating fundamental patterning events, such as the emergence of the AP axis and concomitant germ layer specification, *in vitro*, can therefore help to elucidate body plan and germ layer establishment in *in vivo*.

To tackle this conundrum in a mammalian system, we focus on the first documented symmetry breaking event in gastruloids, which is the polarization of Bra/T expression [14] demarcating initial AP axis establishment [18, 19, 20], and relay our findings to corresponding early lineage specification events in the native mouse embryo. While *symmetry breaking* is a broadly employed term, we hereby refer to asymmetries at the level of cell populations rather than single cells of an aggregate, particularly to the emergence of a Bra/T^+^ posterior domain, prior to which gastruloids do not appear to feature any evident axial identity.

## 3. Results

We employed a Bra/T reporter mESC line (Bra/T::GFP [21]), grown in ESLif (ESL) culture medium, to monitor expression dynamics of Bra/T. Gastruloids were prepared as previously described, including a pulse of (canonical) Wnt agonist CHIR99021 (CHIR99) from 48h to 72h after aggregation (AA) [14, 22] (Figure 1a). In order to quantitatively assess morphology as well as Bra/T expression along the AP axis of the aggregates, a custom made Python-based plugin was used for image segmentation and downstream analysis (Figure 1b).

**Figure 1:**
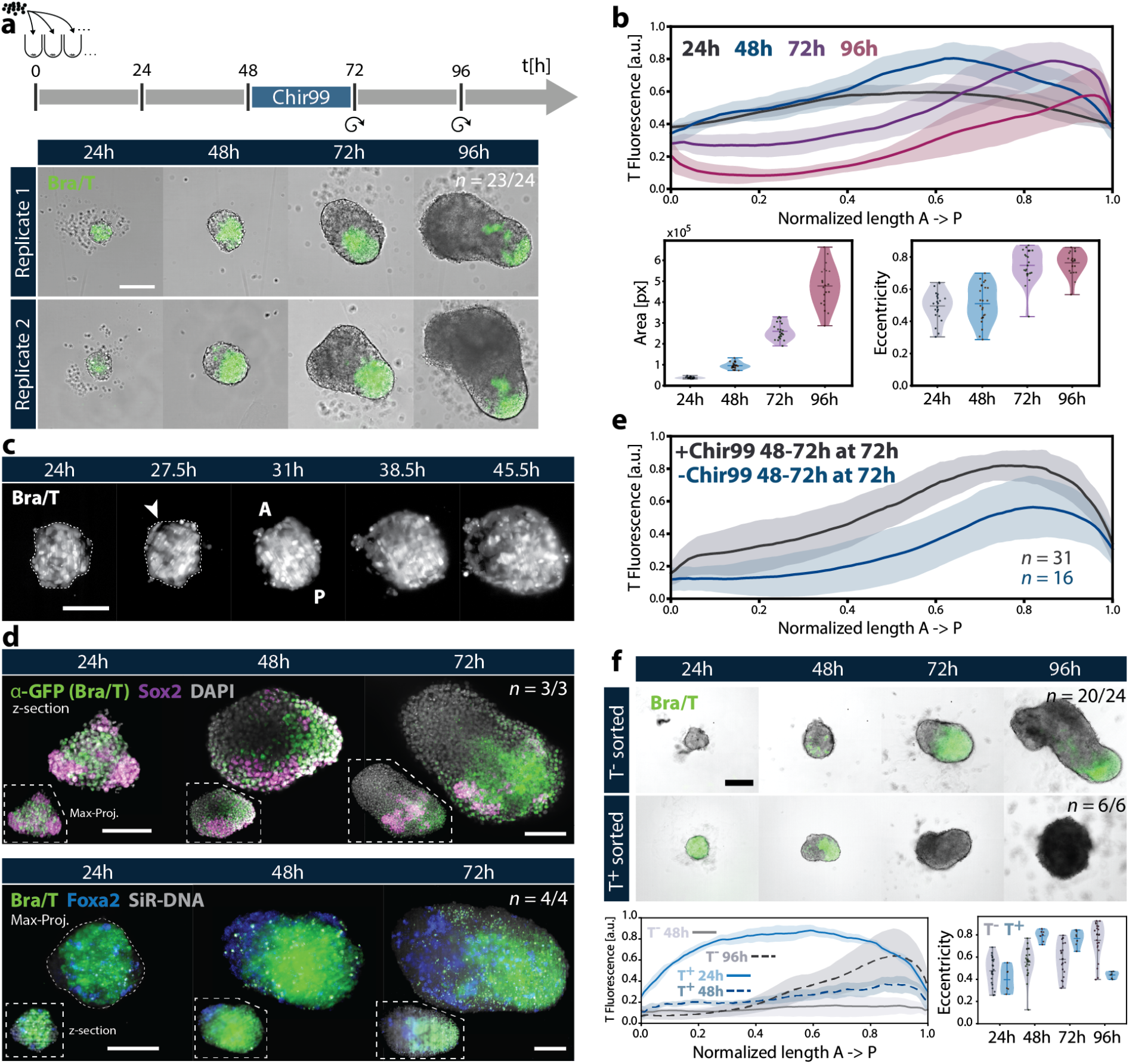
Self-organized AP axis formation in gastruloids demarcated by Bra/T polarization. (a) Gastruloids were generated from Bra::GFP mESCs as previously described [22], including a pulse of Wnt agonist CHIR99 from 48-72h AA. Circular arrows indicate culture medium exchange. Bra/T (T) symmetry breaking occurs around 48h AA and aggregates reach maximum elongation by 96h. Scale bar = 200μm. (b) Quantitative analyses of gastruloids showing Bra/T fluorescence intensity plotted along the normalized AP axis length. Central lines denote the mean intensity values of the included samples, surrounding shades mark the corresponding standard deviation. Shape descriptors, namely area and eccentricity, are also displayed for each timepoint across replicates. (**c**) Light sheet live imaging of Bra::GFP gastruloids to observe the symmetry breaking event. Aggregates at 24h and 27.5h AA are oriented with respect to future AP axis coordinates (from 31h AA onwards). Scale bar = 100μm. (**d**) Depicted are gastruloids around the Bra/T polarization event stained for early germ layer marker genes Sox2 (IHC, pluripotent and neuroectodermal) and Foxa2 (HCR, anterior mesendoderm) together with Bra/T. By 48h, Sox2 forms a domain anteriorly adjacent to Bra/T, Foxa2 is mostly localized to the anterior (i.e. Bra/T^−^) region of the aggregate. There appear to be no clear differences in cell shape between Bra/T^+^ and Bra/T^−^ cells. Scale bars = 100μm. (**d**) Bra/T polarization occurs robustly even without external Wnt activation from 48-72h AA. (**e**) Likewise, gastruloids from both Bra/T^+^ - and Bra/T^−^-sorted Bra/T::GFP mESC cell populations retain the potential to undergo AP axis formation, highlighting their regulative capacity. Scale bar = 200μm.

### 3.1. Emergence of Bra/T polarization

Upon aggregation of mESCs into gastruloids, transcriptional activity of Bra/T rapidly increases within a few hours (Supp. Movie 1). At 24h AA, cells expressing various levels of GFP can be identified throughout the aggregate. By 72h AA, gastruloids across replicates exhibit polarization of Bra/T expression towards the posterior domain and incipient elongation - displayed as an increase in aggregate eccentricity - which proceeds until 96h AA. The exact timing of the symmetry breaking event may vary between experimental batches, but will generally have occurred around or after 48h AA.

We additionally performed light sheet imaging [23, 24] of gastruloids to observe Bra/T polarization at higher resolution (Figure 1c, Supp. Movie 2). It appears that from previously evenly distributed expression, a subtle asymmetry of Bra/T levels along the future AP axis emerges which intensifies as reporter fluorescence further increases in one half of the gastruloid, accompanied by a decreases in the opposite half. This gain and loss of Bra/T expression is concomitant with highly motile behaviour of Bra/T^+^ cells indicating a combination of sorting and signalling that can lead to polarization.

To visualize germ layer formation and their relative spatial arrangement during early gastruloid development in 3D, we performed immunohistochemistry (IHC) for marker genes Sox2 (pluripotent and neuroectoderm) [25] and GFP (mesoderm, via the Bra::GFP reporter) as well as *in situ* hybridization chain reaction (HCR [26]) for Bra/T and Foxa2 (anterior mesendoderm) [27, 28, 29] at developmental timepoints 24h, 48h and 72h (Figure 1d, Supp. Fig. 1a,b).

Imaging of GFP and Bra/T expression through both IHC and HCR in samples between 24h to 72h AA shows that, before and during AP axis specification, Bra/T appears to be (for the most part) homogeneously detectable along the thickness of the aggregate, thereby suggesting that contact- or gravity-dependent Bra/T activation is not required at least for eventual symmetry breaking in gastruloids [11] (Supp. Fig. 1a).

Cells expressing Sox2 are mostly separate from the Bra/T^+^ population with minor overlap. Once the Bra/T pole has emerged, Sox2 forms one or several anteriorly adjacent domains. Notably, there also appear to be Bra/T and Sox2 co-expressing cells at the posteriormost part of the gastruloids, potentially representing NMPs [30].

Foxa2 transcripts are sparsely dispersed throughout the aggregate at 24h AA, prior to Bra/T polarization (Supp. Fig. 1b). Therefore, Foxa2 expression does not seem to be predictive of this event, unlike previous findings in embryoid bodies [11]. After symmetry breaking, Foxa2 is localized opposite of the Bra/T domain towards the anterior of the Gastruloid, in accordance with its role as a marker of the anterior primitive streak.

Additional stainings with phalloidin, demarcating F-actin, in conjunction with IHC for GFP indicate that neither clear epithelial organization nor distinct internal structures are present in early aggregates at 24h as well as polarized gastruloids at 48h AA (Supp. Fig. 1c). By 72h occasional multicellular rosettes have formed in the anterior region (Supp. Fig. 1d). While the core area of the agreggates appears to exhibit elevated F-actin signal across timepoints, there are no clear changes in cell shape or size.

### 3.2. Dispensability of starting conditions for Bra/T polarization

Given that gastruloids often develop a posterior Bra/T domain already by 48h AA, we sought to confirm whether AP axis specification can occur autonomously in gastruloids, i.e. without external Wnt activation and observe that, in gastruloids from Bra::GFP mESCs, Bra/T polarization develops robustly in the absence of the CHIR99 pulse with similar growth and elongation, albeit Bra/T expression is overall weaker by 72h AA (Figure 1d, Supp. Fig. 1d). Likewise, aggregates exhibit similar size and elongation at this timepoint.

Since we observed that ~3% of Bra::GFP mESC grown in ESL display varying levels of GFP expression, gastruloids were generated from either Bra/T^+^ or Bra/T^*−*^ via fluorescence-activated cell sorting (FACS) to thereby probe the necessity of a starting Bra/T^+^ population for symmetry breaking. Strikingly, AP axis formation occurs in both cases, albeit with different developmental timings and dynamics (Figure 1e): In gastruloids from a Bra/T^*−*^ population, dispersed Bra/T^+^ cells emerge starting at 48h AA, which develops into aggregate-wide expression, followed by polarization and elongation by 96h, thereby demonstrating that an initial Bra/T^+^ sub-population is not required for symmetry breaking.

In gastruloids from a pure Bra/T^+^ population, a small subset of Bra/T^+^ cells on one side of the spherical aggregate appears to downregulate Bra/T expression, commencing around 24h, immediately followed by elongation. Thereupon, the Bra*−* domain grows further and Bra/T^+^ cells are only maintained at the elongating tip until 48h AA. Gastruloids subsequently retract to a spherical morphology and disintegrate by 96h AA.

In order to identify whether Bra/T polarization is independent of the signalling environment of the differentiating media, Bra/T::GFP mESCs were seeded in ESL in place of N2B27, the default aggregation medium (Supp. Fig. 1e,f). Also, gastruloids were pulsed with ESL from 0-24h and 24-48h, respectively, thereby facilitating shorter exposure to pluripotency-promoting conditions. We observe that, although Bra/T symmetry breaking is not as reproducible as the control (N2B27) in terms of forming a single pole, GFP^+^ cells nevertheless mostly localize to one side of the aggregate.

Additionally, elongation is impaired in any ESL condition compared to the control such that gastruloids remain mostly spherical by 72h. Altogether this suggests that, while symmetry breaking of Bra/T expression is most robust in differentiation conditions, the molecular mechanisms for developing a distinct Bra/T domain may be dependent on local differences between Bra/T^+^ and Bra/T^*−*^ cell populations which will, however, require future investigation.

### 3.3. Analysis of cell types and populations around the symmetry breaking event

To elucidate the emergence of cell populations and germ layers, concomitant with AP axis specification, we performed single-cell (sc) RNA sequencing of mESCs grown in ESL just prior to aggregation (i.e. trypsinized from 2D culture), representing the 0h timepoint (3812 cells), and of gastruloids at 24h (two replicates, 7480 cells), 48h (two replicates, 9053 cells) as well as 72h (3501 cells) AA.

Resulting datasets were explored via PCA, UMAP clustering, followed by identification of differentially expressed genes as well as GO enrichment analysis (Figure 2a, Supp. Fig. 2a). Based on this and expression of key marker genes, germ layer identities and basic cell states (pluripotent - differentiated) were determined for each cluster.

**Figure 2:**
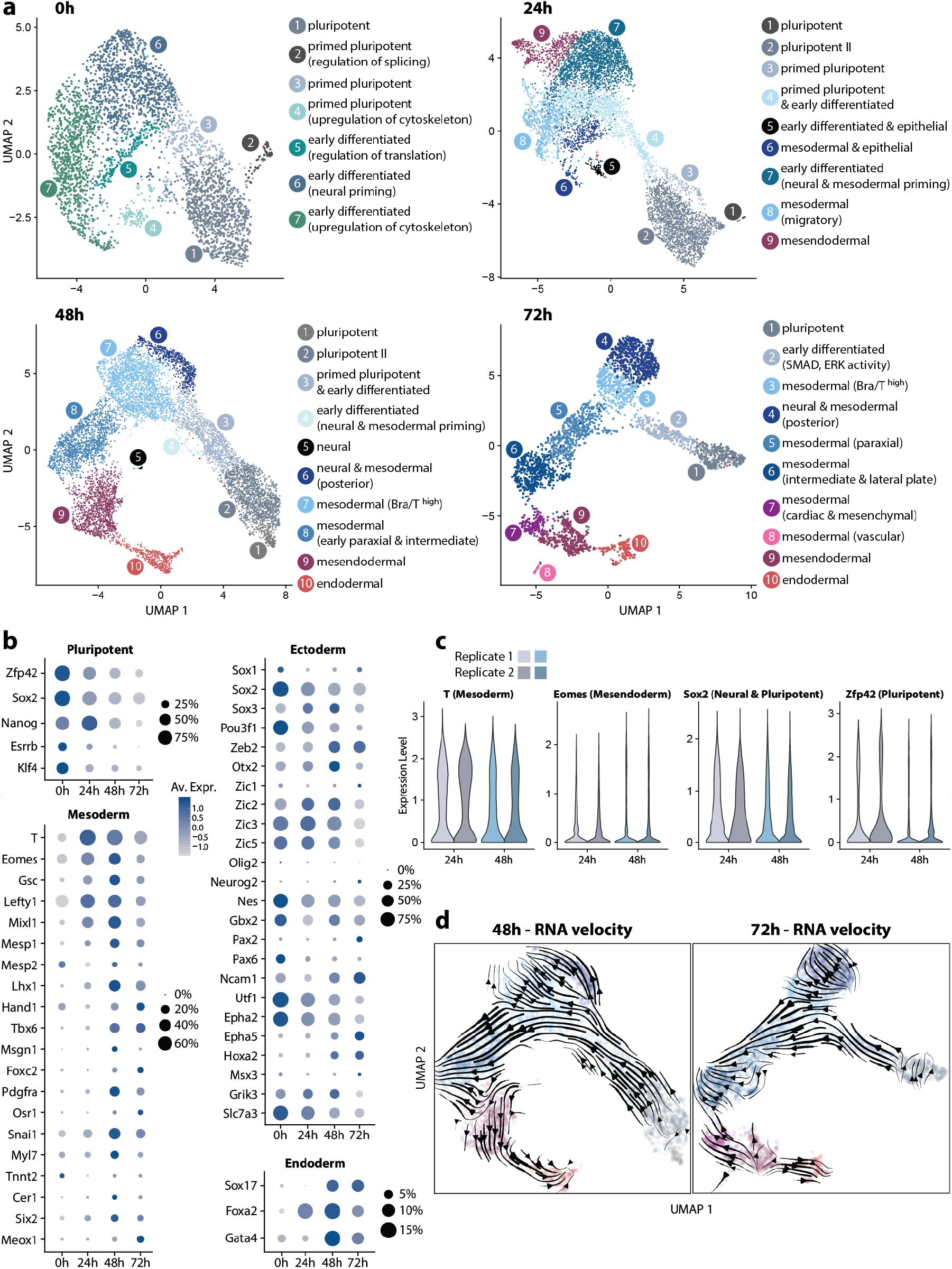
Single-cell transcriptomic analysis of early gastruloid development. (**a**) Germ layer emergence in mESCs (0h) and early gastruloids (24-72h). UMAP plots with clusters annotated according to germ layer or cell state identity are displayed. (**b**) Expression of key marker genes for pluripotent, mesodermal, neuroectodermal or endoderm cells are depicted for each time point. Size of circles denotes fraction of expressing cells and color indicates average scaled expression (Av. Expr.) levels. Note that the size of the circles designates a different fraction in each category. (**c**) Reproducibility of germ layer emergence is demonstrated by separately plotting germ layer marker expression levels of two replicates for 24h and 48h datasets. (**d**) RNA velocity plots on 48h and 72h UMAPs delineate germ layer differentiation trajectories in gastruloids.

At 0h, mostly pluripotent and several primed pluripotent as well as early differentiated cell populations were detected, mostly differing in terms of cytoskeletal, adhesion, splicing and lineage priming gene expression levels. These three populations can also be identified in the later time points which is in contrast to the mouse embryo where epiblast cells by the onset of gastrulation (E6.5), for the most part, do not exhibit naive pluripotency, but rather primed pluripotent or early differentiating states [31, 32]. In gastruloids at 24h, early meso- (Bra/T^+^, Eomes^*−*^) and mesendodermal (Bra/T^+^, Eomes^+^) populations emerge, thereby roughly linking this timepoint to stage E6.5 of the mouse embryo. Distinct neuroectodermal(/neural) populations could not be pinpointed.

Definitive endodermal cells (Sox17^+^), surface at 48h, which is prior to CHIR99 application, hence arguing that, akin to the mesoderm, endoderm lineage can also arise in an autonomous manner. By this time, mesodermal cells have further differentiated into subpopulations (early Bra/T^+^ posterior, paraxial and intermediate). Anterior mesendodermal cells (Eomes^+^) are now largely Bra/T^*−*^ and neural cells are few, the majority of which are mixed with mesodermal cells, without forming a homogeneous cluster. At 72h, lateral plate and cardiac mesoderm-like populations emerge together with a distinct, albeit small cluster of vascular and endothelial progenitors. Notably, a recent study demonstrated how gastruloids can be coaxed into recapitulating early heart morphogenesis by 168h AA via application of cardiogenic factors that are likely enriching these progenitors [33]. We could not identify obvious anterior neural populations, consistent with previous findings in default gastruloids [15].

To further dissect the differentiation dynamics in early gastruloids, we examined levels of germ layer marker genes over time. (Figure 2b). As expected, both the fractions (i.e. percentages) and average expression levels of pluripotent markers decrease, progressing from 0h to 72h with the exception of Nanog that appears to peak at 24h. In regards to the mesoderm, genes Bra/T, Eomes, Lefty1 and Mixl1 are among those which are heavily upregulated at 24h at the onset of germ layer formation in gastruloids. Genes demarcating distinct mesoderm subtypes tend to appear at 48h and 72h (e.g. Pdgfra1, Tbx6, Lhx1, Meox1, Hand1, Mesp1 and 2). This is accompanied by Cadherin-switching, a hallmark of germ layer establishment [34, 35], occurring between 24h-72h AA (Supp. Fig. 2b).

Intriguingly, many early neural lineage markers are highly and pervasively expressed at 0h, which may be explained by the default neural priming of early differentiating cells in standard 2D culture. Expression of some of these neuroectodermal genes thereupon mostly diminishes until 72h (e.g. Nes, Utf1, Epha2, Sox2, Pou3f1), whereas others reach their peak at 48h or 72h (Neurog2, Pax2, Epha5, Hoxa2, Msx3, Ncam1, Zeb2, Zic1). Definitive neural determinant Sox1 is highly expressed in 0h and 72h datasets, although limited to a small subset of cells (*<* 10%). Elevated expression of endodermal markers Sox17 and Gata4 emerges at 48h, congruent with the clustering results. Foxa2, however, surges at 24h, consistent with its role as an early determinant of anterior mesendodermal fate at the onset of gastrulation.

Thereupon, we assessed reproducibility of gastruloids in terms of germ layer generation dynamics by leveraging the replicates available for timepoints 24h and 48h (Figure 2c). Comparing distributions of expression levels of four key marker genes Bra/T (early and posterior mesoderm), Eomes (endoderm, early anterior mesoderm), Sox2 (pluripotent and neural) and Zfp42 (pluripotent) illustrates that, on a population level, gastruloids form germ layers in similar ratios across batches. In addition, performing dataset integration for the respective timepoint yields clear overlap between individual cells from different replicates in both PCA and UMAP plots (Supp. Fig. 2c).

Finally, in order to shed further light on cell differentiation trajectories, we conducted RNA velocity analysis on 48h and 72h datasets (Figure 2d). In both timepoints, there appears to be a distinct progression from pluripotent over primed to early (T+) mesodermal and neural populations. Trajectories towards cardiac mesoderm as well as definitive endoderm on the other hand, originate from a mesendodermal (Eomes^+^) population which is consistent with a recent study that elucidates the roles of Bra/T and Eomes in regulating the exit from pluripotency [36]. Also, among the most pluripotent cells, trajectories orthogonal to the main (mesodermal-heading) flowlines may suggest self-renewal and maintenance of this population.

### 3.4. Size dependence of Bra/T polarization

Following this transcriptional characterization of germ layer emergence, we sought to probe the robustness and plasticity of the symmetry breaking process, amending culture conditions and subjecting gastruloids to a variety of perturbations. Firstly, we varied the initial cell number (*N*) per aggregate between *N* = 30 to 2000 (30, 50, 100, 300, 1000 and 2000) with *N* = 300 as the standard or control condition (Figure 3a, Supp. Fig. 3a,b).

**Figure 3:**
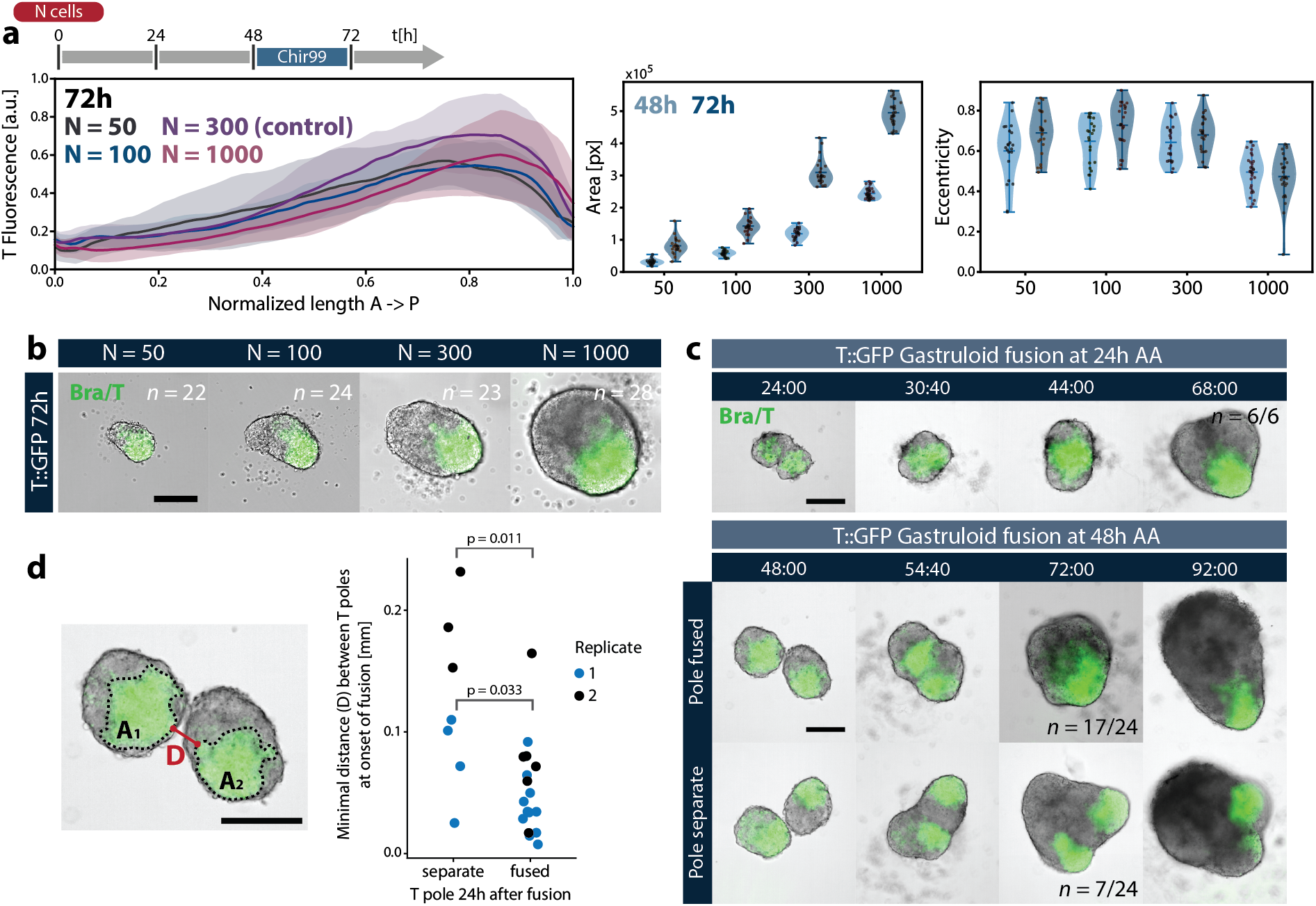
Exploring the regulative capacity of Bra/T polarization in gastruloids. (**a**) Quantitative analyses of gastruloids grown from different initial cell (N) numbers demonstrates reproducibility of Bra/T symmetry breaking and scaling of relative Bra/T domain size between 50-1000 starting cells. Note however how aggregates from N = 1000 do not elongate by 72h as opposed to the other conditions. (**b**) Representative images of gastruloids at 72h AA grown from different N (50,100,300,1000). (**c**) Snapshots of timelapse imaging of aggregates fused together at different timepoints. In case of fusion at 24h AA (prior to Bra/T symmetry breaking), a single Bra/T pole is successfully formed in almost all fusion events, demarcating proper AP axis formation. In case of fusion at 48h AA (AP axis is already formed in individual aggregates), ~29% of fused gastruloids fail to unify their Bra/T domains. (**d**) Successful fusion of Bra/T domains appears to be correlated to the minimal distance D between these at the onset of fusion (timepoint t=0), illustrating the limits of the AP reorganization potential harboured by gastruloids. All scale bars = 200μm.

Notably, Bra/T polarization occurs reproducibly for *N* = 50 − 1000, with gastruloids from different conditions exhibiting scaling of Bra/T domain size along their AP axes. However, aggregates from *N* = 1000 display limited elongation, remaining ovoid by 72h AA, in spite of substantial growth from 48-72h AA (Figure 3a). Using *N* = 2000 yields spherical gastruloids with either no discernable or multiple Bra/T poles, while *N* = 30 results in heterogeneous aggregates, some of which show little to no growth, elongation and Bra/T expression (Supp. Fig. 3a,b). Overall, such a scaling of the Bra/T pole in gastruloids from a wide range of initial cells argues for a self-organizing mechanism that cannot feature a fixed length-scale as in a classical Turing pattern [37].

To further delineate the nature of these interactions, particularly between Bra/T^+^ as well as Bra/T^*−*^ domains and constituent cells, respectively, we performed fusion experiments by placing two gastruloids into the same well of a 96-well plate (96WP) (Figure 3c). When fusing unpolarized samples at 24h AA, we observe that within 48h, i.e. by 96h AA 6/6 develop into a single, larger aggregate with a distinct Bra/T pole. Fusion of gastruloids at 48h AA, when Bra/T symmetry breaking has occurred, yields ~71% (17/24) polarized aggregates at 96h, whereby we observe that Bra/T domains may coalesce and rearrange into one.

Successful pole fusion within 24h appears to be correlated to the distance between the Bra/T domains of the two adjacent gastruloids at the onset of this process (Figure 3d). This suggests that, while Bra/T^+^ cells harbour the propensity to coalesce, a Bra/T^*−*^ region of sufficient size impedes Bra/T^+^ cell migration and/or Bra/T activating signalling cues which is again indicative of the inherent local differences between Bra/T^+^ and Bra/T^*−*^ populations that can lead to polarization.

In order to gauge whether auto-secretion of signaling molecules into the surrounding medium is crucial for Bra/T polarization, we seeded aggregates in 20μl and 200μl of N2B27, instead of the standard 40μl (Supp. Fig. 3c,d). Additionally, for the 200μl condition, medium was exchanged 24h AA. Aggregates grown in 20μl and 200μl generally develop normally and exhibit polarized Bra/T expression by 48-72h AA. Hence, auto-secretion is either robust to media volume alterations and daily exchange or plays a negligible role in gastruloid Bra/T symmetry breaking.

### 3.5. Differences between Bra/T^+^ and Bra/T^−^ populations

Subsequently, we endeavoured to determine how the Bra/T^+^ and Bra/T^*−*^ cells differ from each other in terms of signalling and overall state. Therefore, we leveraged our scRNA datasets and, for each developmental timepoint, separated them into Bra/T^+^ as well as Bra/T^*−*^ populations. Thereupon, differentially expressed features were computed and GO enrichment analysis was performed (Figure 4a-c).

**Figure 4:**
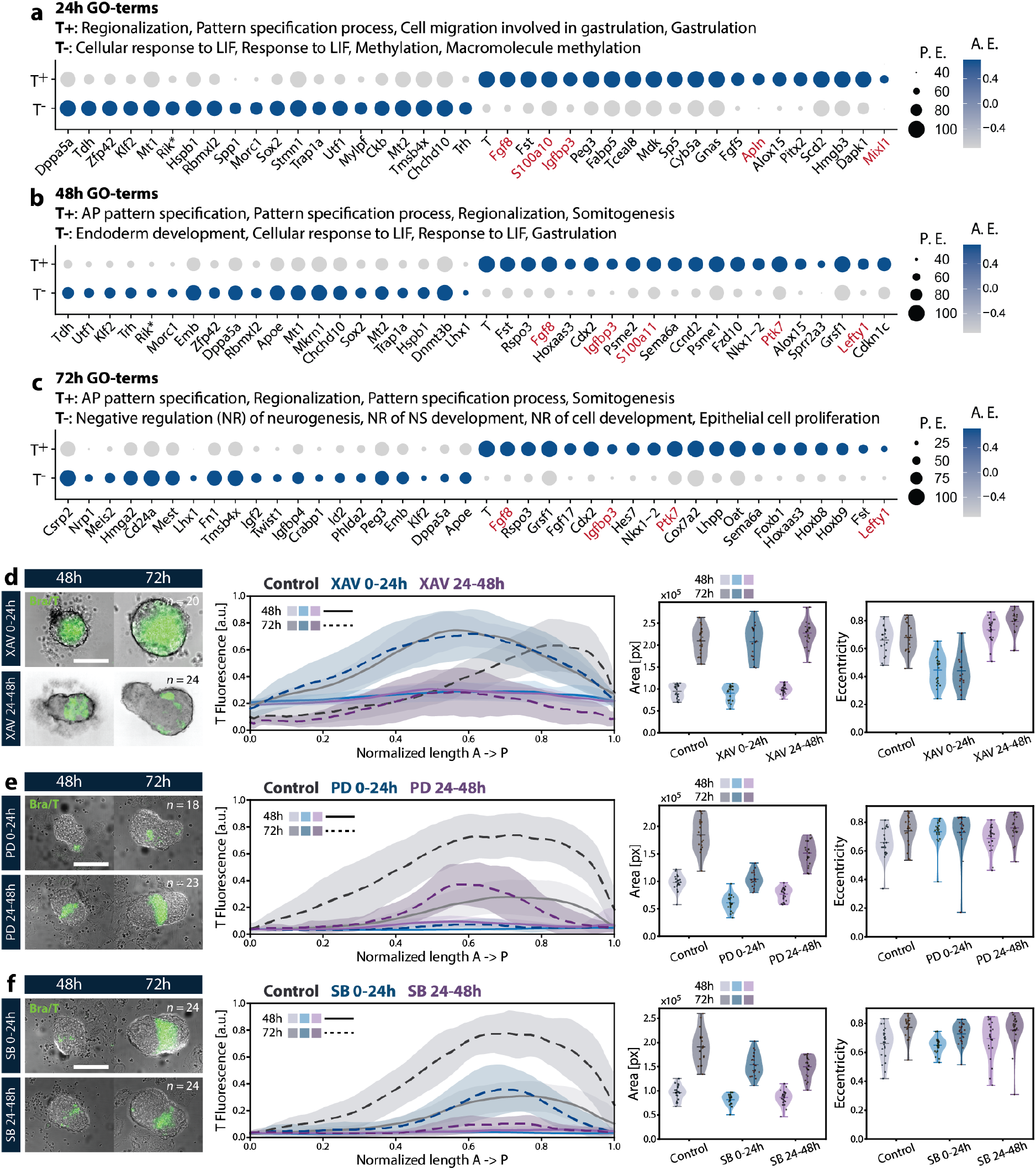
Assessing transcriptional properties of Bra/T^+^ populations and signalling requirements for symmetry breaking. (**a-c**) Single cell transcriptomic comparison of Bra/T^+^ with Bra/T^−^ cell populations around the symmetry breaking event at 48h. Top 4 GO terms as well as top 20 most differentially expressed genes are shown for each population across 3 timepoints (24h, 48h, 72h). For each gene, the size of the circle denotes the fraction of expressing cells (P.E., percentage expressed), while blue colours demarcate elevated (scaled) average expression (A.E.) and grey colours low (scaled) average expression in those cells. Note in particular how Bra/T^+^ populations upregulate genes associated with cell migration and adhesion in the context of EMT and mesoderm formation (highlighted in red). (**d-f**) Timed perturbations of key developmental signalling pathways Wnt via XAV939 (XAV) (**d**), MAPK via PD03 (PD) (**e**) and Nodal/TGF-β via SB43 (SB) (**f**) to dissect their respective roles during Bra/T polarization in gastruloids. Representative images as well as dataset quantifications (Bra/T fluorescence intensity along the gastruloid’s AP axis, area, eccentricity) are shown for each small molecule inhibitor corresponding to a pathway and for each inhibition time frame (0-24h AA and 24-48h AA). Note that the contrasts for the fluorescent channels have been adjusted for display. Scale bars = 200μm.

As expected, throughout all timepoints Bra/T^+^ cells upregulate genes involved in early AP patterning processes, including Wnt, MEK/ERK and TGF-β signalling effectors as well as modulators. This also encompasses somitogenesis-related factors (Fgf8, Hes7) as early as 48h. Moreover, Bra/T^+^ populations feature elevated expression of genes implicated in cell migration, adhesion and epithelial-to-mesenchymal transition (EMT) behaviour such as S100a10, S100a11, Igfbp3, Ptk7, Apln, Lefty1, Mixl1 and Fgf8, a process essential in gastrulation and concomitant mesoderm formation [38].

This is consistent with previous findings [39, 40] and hints at differences in mechanical properties being at the root of the physical separation of the Bra/T^+^ versus the the Bra/T^*−*^ cells in gastruloids. The latter indeed display upregulation of pluripotency as well as early neuroectodermal lineage markers, arguing for distinct cell states between the two populations that consequently entail separate cytoskeletal and adhesive properties which have been shown to underlie sorting of germ layers in other systems [41, 42]. At 48h, Bra/T^*−*^ cells further exhibit elevated expression of endoderm-related genes (Lhx1, Emb) and, by 72h, also of anterior mesodermal and neural determinants (Nrp2, Meis2, Id2, Crabp1).

### 3.6. Developmental signalling during Gastruloid body plan establishment

In order to further study the earliest signalling framework underlying the symmetry breaking process, we dissected the function of key early developmental signalling pathways Wnt, Nodal/TGF-β, MAPK and BMP in Gastruloid AP axis establishment via timed perturbations (Figure 4d-f). Aggregates were thus pulsed between 0-24h AA and 24-48h AA, respectively, with small molecule inhibitors or ligands to disrupt these pathways before or around the time when mesodermal and mesendodermal lineages begin to emerge. At 48h AA, CHIR99 was applied according to the standard protocol, if not stated otherwise.

Inhibition of β-catenin-mediated Wnt signaling by application of XAV939 from 0-24h AA results in a decrease of T transcriptional activity by 48h AA and therefore delays symmetry breaking with aggregates remaining mostly spherical and displaying very little elongation by 72h AA, although Bra/T expression has rebounded and polarization appears to commence (Figure 4d). XAV939 treatment between 24-48h AA does not impede elongation, however Bra/T polarization is perturbed with aggregates at 72h AA mostly exhibiting spotty expression confined to a bulb-shaped, elongation portion of the gastruloid, instead of a continuous Bra/T pole. Aggregate size, however, does not seem to vary substantially between the control and inhibition conditions.

Inhibition of general Wnt secretion by IWP-3 (IWP3) application from 0-24h AA leads to permanent abrogation of symmetry breaking and spherical aggregate morphology with no discernable Bra/T expression (Supp. Fig. 4a). Consequently, treated gastruloids remain much smaller, especially at 72h AA when compared to the control population. Addition of IWP3 between 24-48h AA causes almost identical but slightly less severe effects than treatment between 0-24h AA. Apart from the evident role of canonical and non-canonical Wnts in establishing and maintaining Bra/T expression, this also points towards the latter also playing a key role in axial elongation of gastruloids, as it is known in the mouse embryo, in the context of studies on Wnt5a and Wnt11 [43].

Upregulation of canonical Wnt signaling via pulsing with CHIR99 between 0-24h elicits more robust and clearly restricted Bra/T polarization already at 48h AA and diminished growth, compared to the control condition (Supp. Fig. 4b). Application between 24-48h AA yields spherical, yet Bra/T-polarized gastruloids at 48h AA which thereupon rapidly elongate, appearing to reach slightly higher eccentricities than control aggregates at 72h AA. These results illustrate that, likely due to its inductive effects on Bra/T expression and mesoderm formation, early addition of CHIR99 facilitates and synchronizes Bra/T symmetry breaking and consecutive morphogenetic events, thereby shifting these events to an earlier time point in Gastruloid development.

Inhibition of MEK/ERK signaling through PD0325901 (PD03) at 0-24h AA diminishes Bra/T expression both at 48h and 72h AA (Figure 4e). Whereas elongation is not affected, gastruloids remain smaller and Bra/T-reporter fluorescence is not continuously localized at the posteriormost part, but rather forms a domain or band in the middle or posterior half of the aggregate. PD03 treatment from 24-48h AA yields similar, albeit more striking phenotypes as to reporter expression patterns: Fluorescence intensity levels are higher and Bra/T “band” formation is more pronounced. On the other hand, aggregate growth, while still impeded, does not seem to be as severely affected.

We then perturbed TGF-β signaling in gastruloids using SB-431542 (SB43) and, especially upon inhibition between 0-24h AA, observe formation of a Bra/T domain located in the posterior half, however offset from the elongating tip. This resembles PD03 treated aggregates which, though, generally exhibit a thinner band or stripe of GFP^+^ cells, farther displaced from the posterior end (Figure 4f). We also observe a reduction in gastruloid size at 72h AA in both inhibition conditions.

Notably, for both PD03 24-48hh and SB43 0-24h treatments, this Bra/T “band-like domain” phenotype is still retained upon omitting the 48-72h CHIR99 pulse (data not shown). Live imaging between 48-72h AA of gastruloids pulsed with PD03 at 24-48h and SB43 from 0 to 24h AA, respectively, demonstrates that, in both cases, Bra/T reporter expression emerges in this shape, as opposed to gradual restriction from a continuous domain (Supp. Movies 3-4).

Intriguingly, application SB43 from 0-24h results in larger recovery of Bra/T signal by 72h AA as compared to application between 24-48h AA which is in contrast to PD03 treatment results. This argues that MEK/ERK and TGF-β signalling have at least partly distinct temporal roles in establishment and maintenance or reinforcement of Bra/T expression during early Gastruloid development.

In order to probe whether overactivation of TGF-β signaling affects Bra/T polarization in a similar manner to CHIR99 addition, we applied TGF-β ligand ActivinA (ActA) to aggregates from 0-24h AA as well as 24-48h AA (Supp. Fig. 4c). For both conditions, Bra/T polarization appears to be more clearly restricted to the posterior end of the gastruloid than in the control, both at 48h as well as 72h AA. Although aggregate growth does not seem to be affected, ActA treatment between 24-48h AA further facilitates decreased elongation.

In spite of the implication of BMP signaling in mesoderm formation across species [44, 45], inhibition with DMH1 does not result in a clear effect on gastruloid Bra/T symmetry breaking, elongation and growth, however treated aggregates appear to exhibit aberrant overall and Bra/T domain shapes with higher frequency than the control population, arguing that gastruloid patterning is perturbed to a certain degree (Supp. Fig. 4d).

### 3.7. Comparison of germ layer formation between mouse and gastruloid

While the more differentiated cell types emerging in gastruloids have already been mapped to their respective *in vivo* counterparts as part of previous studies [15, 46, 47, 48], incipient germ layer formation has thus far not been explicitly addressed from such a comparative perspective. Therefore, we conducted mutual nearest neighbor (MNN)-based integration of our gastruloid scRNA-seq dataset from 24h-72h with a previously published mouse sc-atlas [49] (Figure 5a,b).

**Figure 5:**
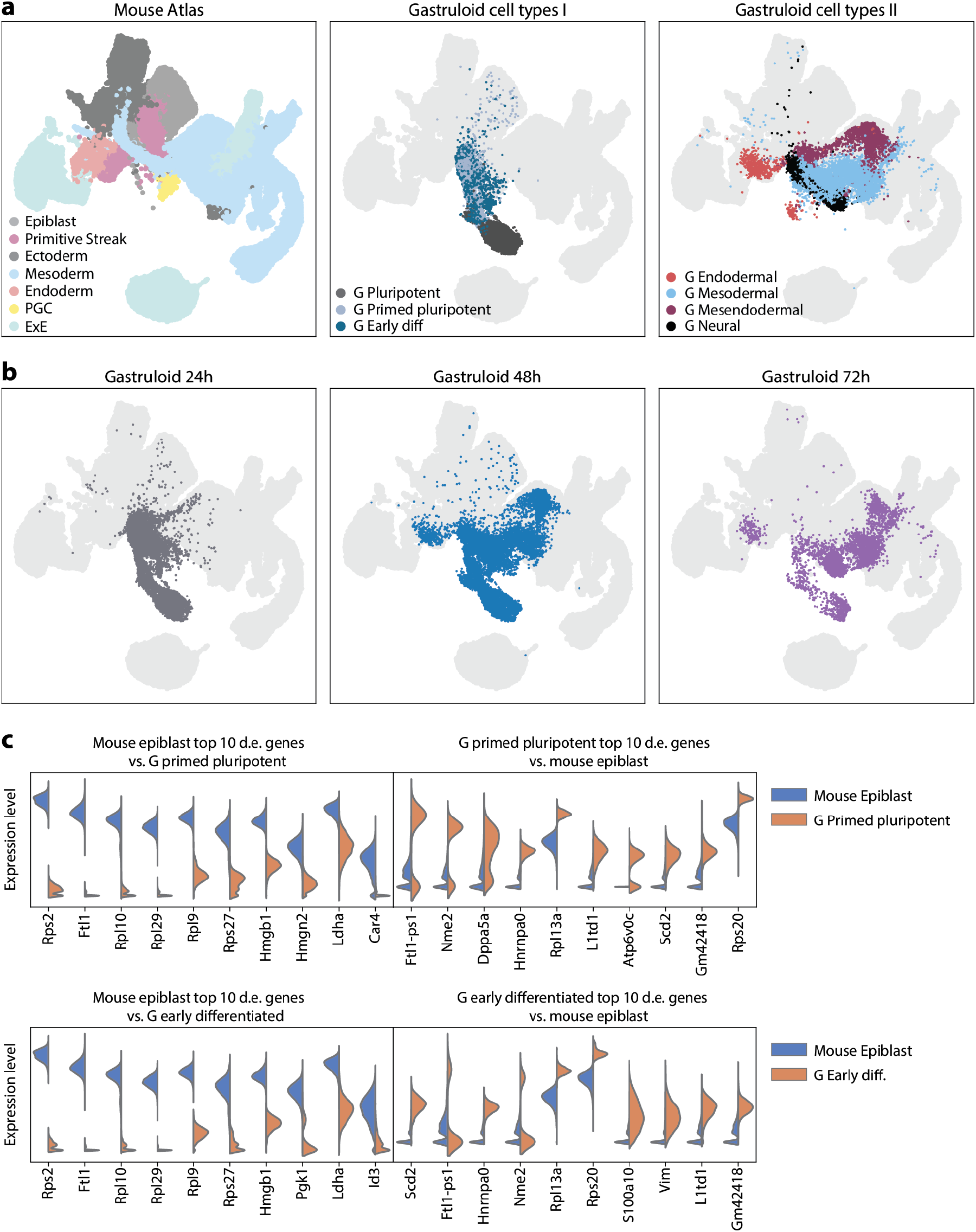
Comparison of cell states and germ layer identities between gastruloids and the early mouse embryo. (**a**) UMAP plots displaying integration of a comprehensive mouse sc-atlas [49] from E6.5-8.5 with the gastruloid datasets of 24h, 48h and 72h. Cell types were labelled according to germ layer identity or differentiation state to identify convergent populations. Note how there is substantial overlap between mesodermal and endodermal cells from both the embryo and the gastruloid, but little intersection between more pluripotent (or epiblast, in case of the mouse) and neuroectodermal populations. (**b**) UMAPs are plotted as above, however, gastruloid cells are filtered by the developmental day instead of germ layer identity or differentiation state. (**c**) Overview of the top 10 most differentially expressed (d.e.) marker genes for the mouse epiblast population versus gastruloid (G) primed pluripotent (pr. pluri.) and early differentiated (e. diff.) populations.

This mouse dataset comprises 9 sequential time-points from E6.5, corresponding to the onset of gastrulation, up to E8.5, characterized (amongst others) by progression of somitogenesis and neural tube closure. Hence, based on clusters found in previous analyses (Figure 2), the gastruloid dataset should roughly correspond to cell populations from E6.5 (24h) to E7.5-8.0 (72h). Germ layer identities were assigned to mouse sc-atlas cell types and to gastruloid UMAP clusters (Figure 2a), in the latter, some clusters feature neuroectodermal-like and mesodermal-like admixture.

UMAP plots of the integrated datasets illustrate that mesodermal and endodermal populations from gastruloids map well onto corresponding cell types. However there is strikingly little overlap between the more undifferentiated populations, including pluripotent, primed pluripotent and early differentiated (i.e. those lacking definite germ layer identity) gastruloid clusters on the one hand, versus the mouse epiblast on the other hand. Importantly, these observations may argue that, in gastruloids, the germ layers form via differentiation trajectories distinct to the mouse embryo, as these seemingly originate from undifferentiated and primed populations featuring a different transcriptional state than the native epiblast.

We additionally observe that, for the most part, neural-like clusters from gastruloids do not seem to map onto neuroectodermal populations in the mouse dataset. This is possibly due to standard gastruloids lacking true neuroepithelia (at least during early developmental stages) while still displaying expression of several neural markers such as Sox1, Sox2, Epha5 and Ncam1.

In order to probe how mouse epiblast cells differ transcriptionally from gastruloid primed-pluripotent and early differentiated populations, the most variable genes were computed (Figure 5c). Epiblast markers thus include ribosome components, metabolic genes such as Car4 (pH regulation, bicarbonate transport) and Ldha (anaerobic glycolysis) as well as chromatin structure modifiers (Hmgb1, Hmgn2). On the other hand, gastruloid primed pluripotent populations display upregulation of distinct translational and metabolic regulator genes, for instance Nme2 (nucleoside triphosphate synthesis), Hnrnpa0 (mRNA metabolism) and Scd2 (fatty acid metabolism), as well as of pluripotency markers Dppa5a and L1td1.

The latter gene remains among the most variable features of the (gastruloid) early differentiated cell population (as compared to the mouse epiblast) which notably further include EMT-associated factors Vim (Vimentin) and S100a10. Conversely, epiblast cells exhibit upregulation of Id3, an epithelial marker gene.

In summary, this argues that, upon exiting pluripotency, gastruloid-forming mESCs immediately adopt a more mesenchymal (non-epithelial) identity by default, comprising a metabolic and cell adhesion state distinct to the mouse epiblast which also helps to explain the comparatively poor mapping of gastruloid neural-like cells to the *in vivo* equivalent. It can therefore be concluded that germ layer formation in gastruloids does seem to occur through developmental trajectories alternative to the mouse embryo.

## 4. Discussion

Studies on embryo-like *in vitro* systems commonly aim to present a faithful replica of native development and hence focus on highlighting similarities to the actual embryo. Here, we characterize symmetry breaking or AP axis establishment in gastruloids and report key differences to the events observed *in vivo* which have not been thoroughly investigated previously. Although embryo-like meso- and endodermal cell types surface during germ layer emergence, neuroectodermal, pluripotent, primed pluripotent and early differentiating populations map comparatively poorly to the mouse epiblast.

The observation that early-primed epiblast-like SCs (EpiLCs) and particularly epiblast SCs (EpiSCs) fail to generate proper, i.e. elongated gastruloids [50] is particularly intriguing in this context. However, it has to be mentioned that a separate study succeeded in generating “Epi-gastruloids”, albeit via comprehensive modifications to the culture conditions [51]. Nevertheless, such results may further corroborate that, using the base gastruloid protocol, AP axis and concomitant germ layer specification cannot efficiently proceed through a direct *in vitro* equivalent of the *in vivo* epiblast cell state.

In addition, the morphogenetic events underpinning symmetry breaking and germ layer formation in gastruloids are also evidently distinct when compared to mouse embryogenesis: Whereas Bra/T^+^ mesoderm always emerges at a specific location from the epithelial layer of the mouse embryo, facilitated by interactions with extraembryonic tissues, in gastruloids, this layer and these tissues are absent. Hence, Bra/T^+^ mesoderm appears to emerge at random positions in the aggregate until a substantial population of Bra/T^+^ cells is generated. Some Bra/T^*−*^ sorted gastruloids may exhibit asymmetric onset of expression which, however does not seem to be correlated to eventual successful pole formation by 96h AA. This, together with the robust symmetry breaking displayed by Bra/T^+^ sorted gastruloids, argues that several initial scenarios can lead to proper AP axis establishment, further highlighting the regulative capacity of mESCs.

To speculate on the regulatory logic of Bra/T emergence and maintenance, it seems that Bra/T expression spreads across cells until 24-48h AA, whereupon the Bra/T^+^ population is roughly maintained until it fades at 96h AA. In conjunction with observations in Bra/T^+^ sorted gastruloids, this may suggest that once a relative threshold of Bra/T^+^ cells have been generated, expression is downregulated in some, likely representing a transition to more differentiated mesoderm subtypes, and further spreading of the Bra/T domain is thereby impeded. It will be worthwhile to further investigate these dynamics and address whether the initial increase in Bra/T^+^ cells is mostly a result of more cell division or of activating signalling.

Following establishment of Bra/T expression dispersed throughout the aggregate, gastruloids proceed to undergo symmetry breaking around or after 48h AA. While the precise mechanisms of this event remain to be elucidated, the comparison of transcriptional profiles of the Bra/T^+^ versus the Bra/T^*−*^ populations in gastruloids performed in this study can provide important clues. Since Bra/T^+^ cells appear to upregulate genes facilitating migratory behavior, it is conceivable that the Bra/T pole is thus formed due to differences in biomechanical properties between Bra/T^+^ and Bra/T^*−*^ populations [42].

Moreover, as mentioned previously, the observed scaling of Bra/T domain size relative to gastruloid AP length in aggregates of different initial cell numbers and sizes, respectively, may argue against a classical Turing-like mechanism underlying the symmetry breaking event. Still, these distinctions will need to be further investigated.

We remark that 2D cell culture conditions of mESCs used to generate gastruloids may substantially influence timing of gastruloid symmetry breaking and germ layer emergence. For instance, gastruloids from mESCs grown in 2i or Serum+2i medium, facilitating a more homogeneous, Nanog-high, naive pluripotent state [52], tend to undergo Bra/T polarization at a later timepoint, usually between 72h and 96h AA [15]. Bra/T::GFPs used in this study were grown in ESL without additional inhibitors and comprise a more heterogeneously pluripotent population, including cells expressing early differentiation markers at low levels (Bra/T, Eomes). Given that a subset of initial cells are already on the threshold of exiting pluripotency, mes(endo)derm in gastruloids emerges rapidly within 24h AA, followed shortly by AP axis formation. This is relevant to note particularly when comparing (e.g. transcriptomic) data from different studies employing distinct cell lines and culture conditions.

### 4.1. An alternative developmental mode?

As demonstrated by studies in basal metazoans, the evolutionarily ancient function of Bra/T appears to be the regulation of folding or invagination movements during Gastrulation [53, 54]. In mammals, Bra/T^+^ cells ingressing at the primitive streak undergo an epithelial-to-mesenchymal transition (EMT) as they form meso- and endoderm [18, 55].

Naturally, this transition facilitates a more migratory state which is at least partly recapitulated in gastruloids. However Bra/T^+^ cells do not ingress from an epithelial layer, which appears to be non-existent at early stages, instead they emerge at seemingly random positions and thereupon arrange into a pole without underlying directional preference, as mentioned above. Likewise, recent studies on endodermal cells in gastruloids have shown that they are specified without the requirement of an EMT as it occurs *in vivo* [56, 57].

Taken together, observations and experimental results elaborated above suggest that germ layer and AP axis formation in gastruloids proceeds via different morphogenetic and transcriptomic developmental trajectories than in the mouse embryo. This is likely an effect of the developmental versatility or regulative capacity of ESCs and ESC-like populations that has been shown to surface in several studied species once such populations are removed from their native embryonic context and grown *in vitro* [58, 59, 60, 61, 62]. For instance, it was recently demonstrated that dissociated and re-aggregated Nematostella gastrulae develop into functional animals which, in contrast to the unperturbed embryo, do not form germ layers via invagination, but rather through delamination, multipolar ingression and cavitation [63].

We propose that the behaviours of such aggregated systems may be summarized as alternative developmental modes which comprise the sum of the (developmental) trajectories of their constituent cell populations. Gastruloids from mESCs thus exhibit an alternative developmental mode which, so far, has not been explicitly characterized in a mammalian model. Furthermore, since gastruloids constitute a system without species-specific external input, that is, absence of native embryonic geometry and extraembryonic tissues, their displayed developmental mode might approximate a more basal one, fundamentally inherent to mESCs [64].

Particularly when considered in conjunction with data from systems that employ extra-embryonic cell types or mimic associated cues, for instance ETS, (i)ETX embryos and blastoids [65, 66, 67, 68], the characterization of such a basal developmental mode as performed in this study can provide deeper insight into fundamental principles of early patterning by helping to delineate which aspects of mammalian body plan establishment are inherent to ESCs and which ones strictly require embryo-like geometry as well as external biochemical and mechanical signals.

Altogether, the analysis of the developmental potential of ESCs aggregated and grown *in vitro* under minimal conditions as conducted in this study can hence shed light on previously uncharted features of early embryogenesis that cannot be gained from studying only the native embryo.

## 5. Material & Methods

### 5.1. mESC culture

mESCs were cultured as previously described [22]. In brief, Bra::GFP mESCs were maintained in ES-Lif (ESL), consisting of KnockOut D-MEM supplemented with 10% fetal bovine serum (FBS), 1x Non-essential amino acids (NEEA), 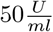 Pen/Strep, 1x GlutaMax, 1x Sodium Pyruvate, 50μM 2-Mercaptoethanol and leukemia inhibitory factor (LIF, homemade), on 0.1% gelatin-coated (Millipore, ES-006-B) tissue culture-treated 25cm^2^ flasks (T25 flasks, Corning, 353108) in a humidified incubator (37°C, 5% CO_2_). ESL was prepared and provided by the Tissue Engineering Unit of the Centre for Genomic Regulation (CRG). mESCs were passaged every 2-3 days and the ESL was exchanged for fresh, prewarmed medium on alternating days. Cells were cultured up to 50-70% prior to passaging or experimental use.

### 5.2. Generation of gastruloids

Gastruloids were generated as previously described [22]. To sum up, mESCs growing in ESL were gently rinsed with 5ml pre-warmed DPBS^+*/*+^ (DPBS with Mg and CaCl_2_, Sigma, D8662). DPBS^+*/*+^ was replaced with 1ml pre-warmed Trypsin-EDTA (Gibco, 25300) and the cells were incubated at 37°C for 1-2min. Thereupon, Trypsin-EDTA was neutralized via application of 4ml ESL. Cells were collected via gentle pipetting, the suspension was transferred to a 15ml centrifuge tube and centrifuged for 3min at about 180g (900-1000rpm).

Then the supernatant was aspirated and cells were resuspended in 10ml warm DPBS^+*/*+^ (washing). This step was repeated and after another centrifugation round (3min, 180*g*), the pellet was resuspended in 0.5ml to 1.5ml of pre-warmed differentiation medium N2B27 (Ndiff227, Takara, Y40002) via pipetting up-and-down 5-15 times with a P1000 pipette until remaining clumps of mESCs were separated into single cells.

Hereafter, mESCs were counted using an automated cell counter (Countess II, Invitrogen). For this, 10μl of cell suspension were mixed with 10μl staining mix (Invitrogen, T10282) and loaded on a counting slide (Invitrogen, C10228). The calculated volume of the cell suspension was then added to the required amount of warm N2B27 and transferred to a sterile reservoir. Using a microchannel pipette, 40μl of plating suspension were added to each well of a U-bottom, low adhesion 96-well-plate (96WP, Greiner, 650970) which was thereafter returned to the incubator and maintained at 37°C and 5% CO_2_.

After 48h, a further 150μl of prewarmed N2B27 containing 3μM CHIR99 (Sigma, SML1046) were pipetted into each well, followed by daily medium exchange with just N2B27 if gastruloids were grown beyond 72h.

### 5.3. Application of small molecules and ESL pulse

In case of application from 0-24h, small molecules were diluted to the desired concentration in warm N2B27 just prior to addition of the counted cell suspension. Gastruloids were then plated, using this mixture, as described above. The next day, small molecules were diluted via addition of 150μl N2B27 and subsequent removal (washing) after 5min to allow the aggregates to sink to the bottom of the 96WP.

In case of application from 24-48h, 24h after plating, 20μl of N2B27 per Gastruloid were replaced with 20μl of warm N2B27 containing the respective small molecule diluted to 2x the final desired concentration. 48h AA, gastruloids were washed once with 150μl N2B27, thereupon 3μM CHIR99 were applied in 150μl.

Applying ESL to gastruloids from 0-24h was performed by diluting counted mESCs into pre-warmed ESL, instead of N2B27, followed by plating of gastruloids. At 24h AA, gastruloids were washed twice with 150μl and thus left in 40μl media volume until the CHIR99 pulse. For application from 24-48h, gastruloids were washed twice with 150μl ESL and left in 40μl medium. At 48h AA, prior to applying 3μM CHIR99 in 150μl N2B27, gastruloids were again washed twice with 150μl N2B27. Table 1 provides an overview of small molecules employed in this study and corresponding concentrations.

**Table 1:** List of small molecules with dilutions

| Small molecule | Dilution (stock to final) | Stock diluted in | Source |
| --- | --- | --- | --- |
| CHIR99 | 10mM to 3μM | DMSO | Sigma, SML1046 |
| XAV939 | 10mM to 1μM | DMSO | Sigma, X3004 |
| IWP3 | 5mM to 1μM | DMSO | Sigma, SML0533 |
| PD03 | 10mM to 1μM | DMSO | Sigma, PZ0162 |
| SB43 | 10mM to 10μM | DMSO | Sigma, S4317 |
| ActA | 100 $\frac{\mu g}{ml}$ to 50 $\frac{ng}{ml}$ | 2mM HCl | R&D Systems, 338-AC-010 |
| DMH1 | 10mM to 1μM | DMSO | Sigma, D8946 |

### 5.4. Immunohistochemistry

For fixation, aggregates were harvested using organoid collection plates (OCPs, patent EP3404092A1) and subsequently pooled in 2ml Eppendorf tubes with a cut-off P1000 pipette tip. After 3 washes for 5min each with 1ml DPBS_*−/−*_ (DPBS without Mg and CaCl_2_, Sigma, D8537), 1ml 4% paraformaldehyde (PFA) in DPBS_*−/−*_ was applied and samples were incubated for 2h at 4°C with gentle horizontal rotation. Hereafter, gastruloids were washed 3-5x with 1ml DPBS−/− each. At this point, aggregates were optionally stored at 4°C for a few days to 1 week.

All subsequent steps were conducted at 4°C and under gentle shaking, if not stated otherwise: Aggregates were washed 3x for 10min with 500ul DPBS_*−/−*_ containing 9% foetal bovine serum (FBS), 1% BSA (Jackson Immuno Research, 001-000-162) and 0.2% Triton X-100 (PBSFBT). For blocking, gastruloids were incubated for 2h in PBSFT on an orbital shaker. In case of single-round antibody (AB) stainings, anti-GFP-AF647 (ThermoFisher, A-31852) and Phalloidin-AF555 (ThermoFisher, A34055) were applied at 1:500 in 500ul PBSFBT overnight (ON).

The next day, samples were washed with PBSFBT 3x for 30min. Afterwards, aggregates were washed 3x with DPBS_*−/−*_ containing 0.2% FBS and 0.2% Tween (PBT) for 15min each. Gastruloids were then removed from the orbital shaker and PBT was replaced with 100μl Vectashield with DAPI (Vector Laboratories, H-1200-10) per 2ml tube. Using a cut-off P200 tip, 10-20 gastruloids per condition were then transferred into a well of a flat-bottom CellCarrier-96 ultra plate (PerkinElmer, 6007008) and imaged on an Opera Phenix HCS system (PerkinElmer) in the confocal mode, using a 20x water objective.

In case IHC was performed with primary and secondary ABs, primary ABs chicken anti-GFP (Sigma-Aldrich, AB16901) and rabbit anti-Sox2 (ThermoFisher, PA1-094) were applied at 1:250 in 500ul PBSFBT overnight (ON). The following day, samples were washed 6x with PBSFBT: 3x for 10min and 3x for 30min. At this point, samples can be kept ON. Subsequently, secondary ABs goat anti-chicken AF488 (Invitrogen, A-11039) and donkey anti-rabbit AF568 (Invitrogen, A10042) were applied, diluted 1:250 in 500ul PBSFBT either ON or for 2h at 32°C in a heating block (ThermoMixer C, Eppendorf) with integrated shaking (350rpm). Then, aggregates were washed 7x: 3x for 10min and 4x for 30min. Hereafter, 3 more washes were performed with PBT for 15min each. Finally, PBT was replaced with 100μl Vectashield with DAPI (Vector Laboratories, H-1200-10) per 2ml tube. Samples were then mounted and imaged as described above.

### 5.5. *In situ* HCR

Gastruloids were harvested from round bottom 96WPs in UV-sterilized OCPs and transferred to 2ml Eppendorf tubes using RNase free wide-bore P1000 pipette tips. After 3 washes for 5min each with 1ml DPBS_*−/−*_, samples were fixed ON at 4°C in 1ml 4% PFA in DPBS_*−/−*_. The following day, samples were dehydrated by 3 DPBS_*−/−*_ washes (5min each, room temperature), 4 washes in 100% Methanol (MeOH) for 10min each at room temperature (RT) and 1 wash in 100% MeOH for 50min at RT. Hereafter, samples were stored at −2-°C, if required.

Rehydration was performed through a series of sequential washes at RT: 5min in 75% MeOH in DPBS_*−/−*_ with 0.1% Tween (PBST2), 5min in 50% MeOH in PBST2, 5min in 25% MeOH in PBST2 and 5x in PBST2 for 5min each. Samples were then transferred to a 1.5ml Eppendorf tube and 500μl probe hybridization buffer (PHB, Molecular Instruments, HCR v3.0) was applied for 30min at 37°C. Immediately afterwards, probe solution was prepared by adding 2pmol of each probe of interest (Foxa2 and Bra/T, both ordered through Molecular Instruments for HCR v3.0) into 500μl PHB and maintained at 37°C for 30min. Thereupon, probe solution was applied to the samples, replacing the PHB and incubation was performed ON at 37°C.

The next day, 4 washes (15min each, 37°C) were conducted in 500μl probe wash buffer (PWB, Molecular Instruments, HCR v3.0) preheated to 37°C and 2 washes (5min each, RT) were performed in 5x SSCT (5x sodium chloride sodium citrate, 0.1% Tween in ultrapure H_2_O). Hereafter, amplification buffer (Molecular Instruments, HCR v3.0) was equilibrated at RT and SSCT was replaced with 500μl amplification buffer (equilibrated at RT), followed by incubation for 30min at RT.

30pmol of each hairpin (h1 and h2m, Molecular Instruments, HCR v3.0) were then separately aliquoted (2μl of 3μM stock), heated to 95°C for 90s and cooled for 30min in the dark at RT. Thereupon, hairpins were mixed in 500μl amplification buffer, pre-equilibrated at RT, to a final concentration of 60nM per hairpin. Amplification buffer was removed from samples and replaced with hairpins in amplification buffer, followed by incubation ON at RT, in the dark.

Hereafter, 5 washes in SSCT were performed at RT: 2x 5min, 2x 30min and 1x 5min. Nuclear stain SiR-DNA (Spirochrome) was added with the final wash at a dilution of 1:1000. Samples were then stored at 4°C in the dark for 24h to 2 weeks prior to imaging on a MuViSpim CS (Bruker).

### 5.6. Imaging and quantitative analysis of gastruloids

Cell aggregates cultured in round-bottom 96WPs were imaged with an Opera Phenix HSC system (PerkinElmer) in wide-field mode using a 10X air objective (NA 0.3). Focus heights were manually adjusted for a given plate, depending on size of the gastruloids. For time lapse experiments, the incubator module was set at 37°C and 5% CO_2_. Medium evaporation was prevented by replacing the lid of the round-bottom 96WP with a MicroClime environmental microplate lid (Labcyte, LLS-0310) filled up with DPBS_*−/−*_ to maintain humidity. Images were then acquired with a 10 mins interval.

After acquisition, images were compiled in single multichannel files. All images were segmented via a machine learning (ML) pipeline implemented with a custom-written Python code. The segmentation was performed using only the bright-field (BF) image: 2 to 3 images per plate (~3%) were randomly chosen and a mask was manually annotated, thus generating the ground truth (GT) images. BF and GT were then used as input of the downstream ML pipeline. Firstly, images were downsampled by a factor of two to improve computation speed.

Next, for all input BF images, approximately 350 features were extracted, including difference of gaussians, gradients of gaussians, Laplace filters and DAISY descriptors with different variance values. A logistic regression model was trained to predict, for each pixel, the probability of being classified as background, inside gastruloid, or gastruloid edge. The trained model was thereupon used to classify the pixels in all other images in the 96WP.

Because of the variability observed in BF image quality an additional step was implemented to ensure proper segmentation of the gastruloids. The classification output generated an image containing the identity of every pixel as well as the probability map for the “gastruloid edge” class. Using the second map, watershed segmentation was used to generate an alternative segmentation. As a result of this preliminary segmentation, we hence obtained: i) a classifier mask and ii) a watershed-based mask.

Upon visual inspection, we observed that the watershed mask was accurate enough and properly segmented the gastruloid image in approximately 90-95% of the cases. Otherwise, the classifier mask could be employed to yield an accurate segmentation. For the few cases in which neither watershed not classifier masks produced an acceptable segmentation, the pipeline further includes the option to generate a mask manually.

The final mask was smoothened using standard morphological operations, and the area as well as the perimeter of the gastruloid were extracted. To account for the bending of gastruloids during elongation, eccentricity was computed on a computationally straightened version of the final mask. Briefly, we used distance transform to find the midline and the width of the gastruloid, and computed eccentricity according to 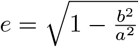, here a and b represent the major and minor axis length of the ellipse with the same second moments as the straightened gastruloid mask.

To compute the AP fluorescence intensity profiles, for every position along the midline of the gastruloid, the average pixel intensity in the fluorescence image channel along the direction orthogonal to the midline was computed. The AP profile was then oriented such that the posterior part of the gastruloid always represented the highest Bra/T expressing pole. For analysis across conditions, the gastruloid length was normalized to values between 0 (anterior) and 1 (posterior).

### 5.7. Fusion experiments and data analysis

In order to perform fusion experiments of gastruloids at desired timepoints, single aggregates grown in round-bottom 96WPs were extracted from their respective well by pipetting up-and-down twice with cut-off P200 tip. The entire media volume (40μl), including the Gastruloid, was then released into a designated well already containing another aggregate. Time-lapse imaging was performed as described above on an Opera Phenix HCS system (PerkinElmer).

For distance analysis between the Bra/T domains of the fusing aggregates, domains were segmented in Fiji/ImageJ [69, 70], using the “Threshold” function (Default and Dark background were selected). “XStart” and “YStart” values of the thus binarized masks were then determined via the “Analyze Particles” function (Display results, Record starts and Include holes were selected). Hereafter, the “Graph” plugin was employed to determine the minimal distance (D) between the two segmented Bra/T domains. Results were exported as csv files. Statistical tests were performed and results were plotted in R version 4.0.3.

### 5.8. Preparation of gastruloids for scRNA-seq

To ensure sufficient numbers of dissociated cells for each experimental condition, 5 96WPs of gastruloids were generated for the 24h timepoint, 3 for the 48h timepoint and 2 for the 72h timepoint. For the 0h timepoint, Bra/T::GFP mESC were grown in a T25 flask as described above. Individual plates were harvested using UV-sterilized OCPs and gastruloids were pooled into a 15ml centrifuge tube using RNase-free wide-bore P1000 pipette tips (Thermo Fisher Scientific, 2079G). When all plates from a given condition were harvested, the centrifuge tube was placed into an incubator (37°C, 5% CO_2_) and harvesting proceeded with plates from the next condition.

Once all samples were collected, thus yielding one 15ml centrifuge tube per condition, aggregates were centrifuged at 180*g* for 2min. Hereafter, the supernatant was removed and 1ml TrypLE (Thermo Fisher Scientific, 12604) was added per tube for dissociation and samples were incubated for 5min at (37°C, 5% CO_2_). Then, TrypLE was deactivated using 4ml ESL and gastruloids were centrifuged for 2min at 180*g*. Supernatant was removed, samples were washed with 5ml cold (4°C) 0.1% RNase-free BSA (Thermo Fisher Scientific, AM2616) in DPBS_*−/−*_ (BPBS) and centrifuged (2min, 180*g*). From this step onwards, samples were maintained at 4°C.

Following removal of the supernatant, the pellet was resuspended in 50-100μl cold BPBS (depending on pellet size) by gently pipetting up and down 3-5x with a P200 tip. The cell suspension was then filtered through a 35-μm filtered cap tube (Corning, 352235). Cells were counted as previously described, aiming for a concentration of around 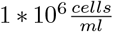. In case of high concentrations 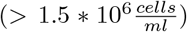, the cell suspension was diluting with cold BPBS and re-counted.

For the 0h timepoint, mESCs were trypsinized using 1ml Trypsin-EDTA 0.05% as described above. Following neutralization (4ml ESL) and centrifugation (180g, 3min), the supernatant was discarded and cells were washed with 5ml cold BPBS. Cells were then kept on ice, until samples from other experimental conditions reached this washing step. At this point, all samples were joined together for centrifugation and further procedures illustrated above.

### 5.9. Library preparation and sequencing

After the generation of single-cell suspensions in BPBS as described above, cells in each sample were barcoded with a Chromium Single Cell 3’ GEM, a Library & Gel Bead Kit v3 (10x Genomics) and a Chromium Single Cell B Chip Kit (10x Genomics) on a Chromium Controller (10x Genomics), followed by cDNA library construction according to manufacturer’s instructions. These libraries were sequenced via a NextSeq 500 system (Illumina). We read 8, 28, and 56-base pairs for Tru-seq indices, 10x barcodes with unique molecular identifiers (UMIs), and fragmented cDNA, respectively.

### 5.10. Basic scRNA-seq data analysis

Quality control (QC), alignment to the mouse genome (GRCm38), and counting of the sequence reads were conducted with CellRanger v3.1.0 (10x Genomics) to generate feature-barcode matrices. The summary of the statistics of sequencing results is shown in supplementary table X.

Further QC steps, normalization, identification of most variable features, scaling, PCA and clustering were performed with Seurat version 3.2.3 loaded into R version 4.0.3. Low quality cells were removed with the following thresholds: Unique feature counts (UFC) *>* 2500 & mitochondrial counts (MC) *<* 10% & Total RNA counts (TRC) *<* 150000 for the 0h dataset, UFC *>* 2750 & MC *<* 20% & TRC *<* 100000 for the 1st replicate of the 24h dataset, UFC *>* 3750 & MC *<* 15% & TRC *<* 150000 for the 2nd replicate of the 24h dataset, UFC *>* 2750 & MC *<* 20% & TRC *<* 100000 for the 1st replicate of the 48h dataset, UFC *>* 3500 & MC *<* 15% & TRC *<* 120000 for the 2nd replicate of the 48h dataset and UFC *>* 2500 & MC *<* 20% & TRC *<* 135000 for the 72h dataset.

24h and 48h datasets were integrated using the “FindIntegrationAnchors()” and “IntegrateData()” functions. Following assessment of cell cycle bias on clustering results via the “CellCycleScoring()” function which is based on expression G2M and S-phase marker genes, cell cycle correction was peformed on the 0h and 24h datasets and rescaled data was reclustered and re-plotted.

Most differentially expressed cluster marker genes were identified through the “FindAllMarkers(…, min.pct = 0.25, logfc.threshold = 0.25)” function. GO-term analyses were performed using “ClusterProfiler”. For comparing expression of selected germ layer genes between all timepoints, datasets were merged with the “merge()” function.

In order to analyze and compare Bra/T^+^ with Bra/T^*−*^ populations across timepoints, cells from each dataset were separated based on a normalized expression level cut-off of 0.5, with *>*0.5 = Bra/T^+^ and *≤*0.5 = Bra/T^*−*^.

### 5.11. RNA velocity analysis

The scRNA sequencing dataset of gastruloid cells from 48 and 72h AA were used to perform RNA velocity analysis. To this aim, we used one replicate and first generated an annotated loom file using the velocyto Python package [71] and the genome annotation file from GENCODE. Next, RNA velocity was performed using scVelo in ?stochastic? mode [72]. The resulting embedding streams were overlaid on the UMAP plot and cells were color-coded according to their cluster identity.

### 5.12. Integration of gastruloid and mouse scRNA datasets

Gastruloids scRNAseq data obtained from 24, 48 and 72h AA were integrated to a comprehensive reference dataset [49] using mutual nearest neighbor as previously described [73]. Briefly, datasets were normalized and the counts of the highly variable genes were used for principal component analysis (number of PCs=50). All the downstream analysis was performed in PC space. First, batch correction was performed on the concatenated datasets using the R function “reducedMNN” from the ?batchelor? package. The corrected dataset of the 50 PCs was then used as input of the “umap” R function (*n*_*n*_*eighbors* = 20, *min*_*d*_*ist* = 0.7). Cell clusters from mouse and gastruloids were redefined to consider the major germ layer lineages and individual cells were plotted on the common UMAP space and color-coded according to cluster identity or developmental stage.

Differential expression analysis was performed considering pairs of cell clusters. Counts were log-normalized and the average expression within each cluster was computed. Next, highly differentially expressed genes were identified using a differential cut-off of +− 1. In the main figures, only the top10 differentially expressed genes are represented.

## 7. Acknowledgments

The Bra::GFP cell line is a kind gift from Alfonso Martinez Arias. We thank the Universitat Pompeu Fabra (UPF) genomics core facility and EMBL Heidelberg GeneCore for the sequencing, the Mesoscopic Imaging Facility at EMBL Barcelona for assistance with imaging, the CRG/UPF FACS Unit for help with sorting experiments as well as members of the Vikas Trivedi and Miki Ebisuya labs at EMBL Barcelona for discussions and critical feedback. We further thank Blanca Pijuan-Sala, Leah Rosen and Carine Stapel for feedback on the scRNA-seq analysis as well as Ricard Argelaguet for advice on and providing code for scRNA dataset integration. This work was supported by funds from the European Molecular Biology Laboratory; NG and FN also acknowledge the Human Frontier Science Program (HFSP) for financial support. DO was supported by a Juan de la Cierva Postdoctoral Fellow-ship.

