## Supplementary Material for "Dynamics of anteroposterior axis establishment in a mammalian embryo-like system"

[*]These authors contributed equally.
[1]EMBL Barcelona, 08003 Barcelona, Spain.
[2]Institució Catalana de Recerca i Estudis Avançats, 08010, Barcelona, Spain.
[3]EMBL Heidelberg, Developmental Biology Unit, 69117 Heidelberg, Germany.

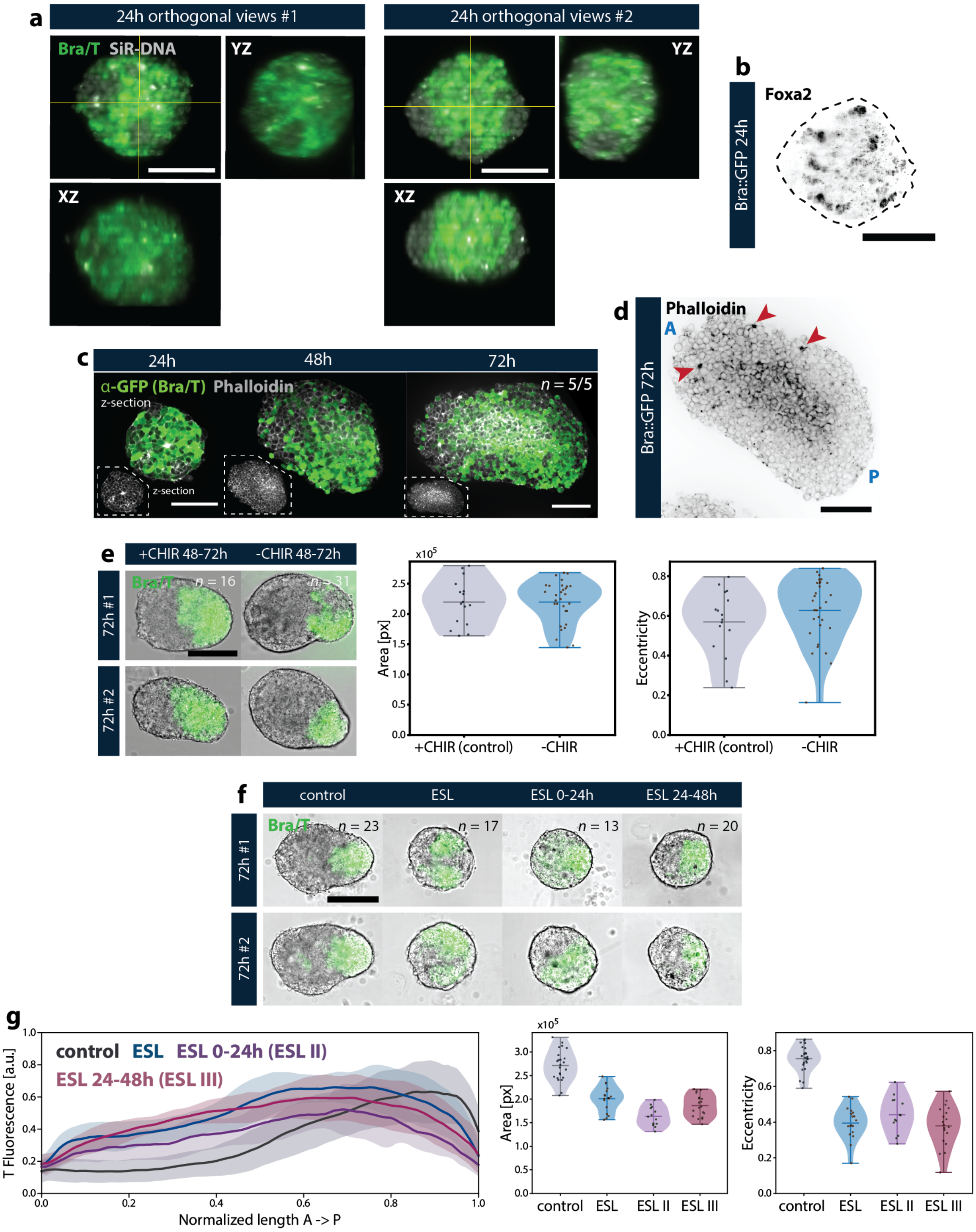


Supplementary Figure 1: **Self-organized AP axis formation in gastruloids demarcated by Bra/T polarization**

*(**a**) IHC for Bra/T in gastruloids at 24h AA with orthogonal views demonstrates that expression is evenly distributed along the thickness of the aggregate. Scale bars = 100µm. (**b**) HCR for Foxa2 illustrates dispersed, but evenly distributed expression in gastruloids at 24h AA, prior to Bra/T symmetry breaking. Scale bar = 100µm. (**c,d**) IHC for Phalloidin in gastruloids at 24h, 48h and 72h AA indicates no clear differences in cell shape between Bra/T$^{+}$ and Bra/T$^{-}$ cells. However, there is occasional multicellular rosette formation in the anterior (A) region (red arrows) of gastruloids at 72h AA. Scale bars = 100µm. (**e**) Representative images for gastruloids grown with or without canonical Wnt upregulation (CHIR99 pulse) as well as quantifications of aggregate area and eccentricity (or elongation). Scale bar = 200µm. (**f**) and (**g**) Representative images for gastruloids grown under pluripotency-promoting conditions via ESL and corresponding quantifications of Bra/T fluorescence intensity along the AP axis as well as aggregate area and eccentricity. Scale bar = 200µm.*

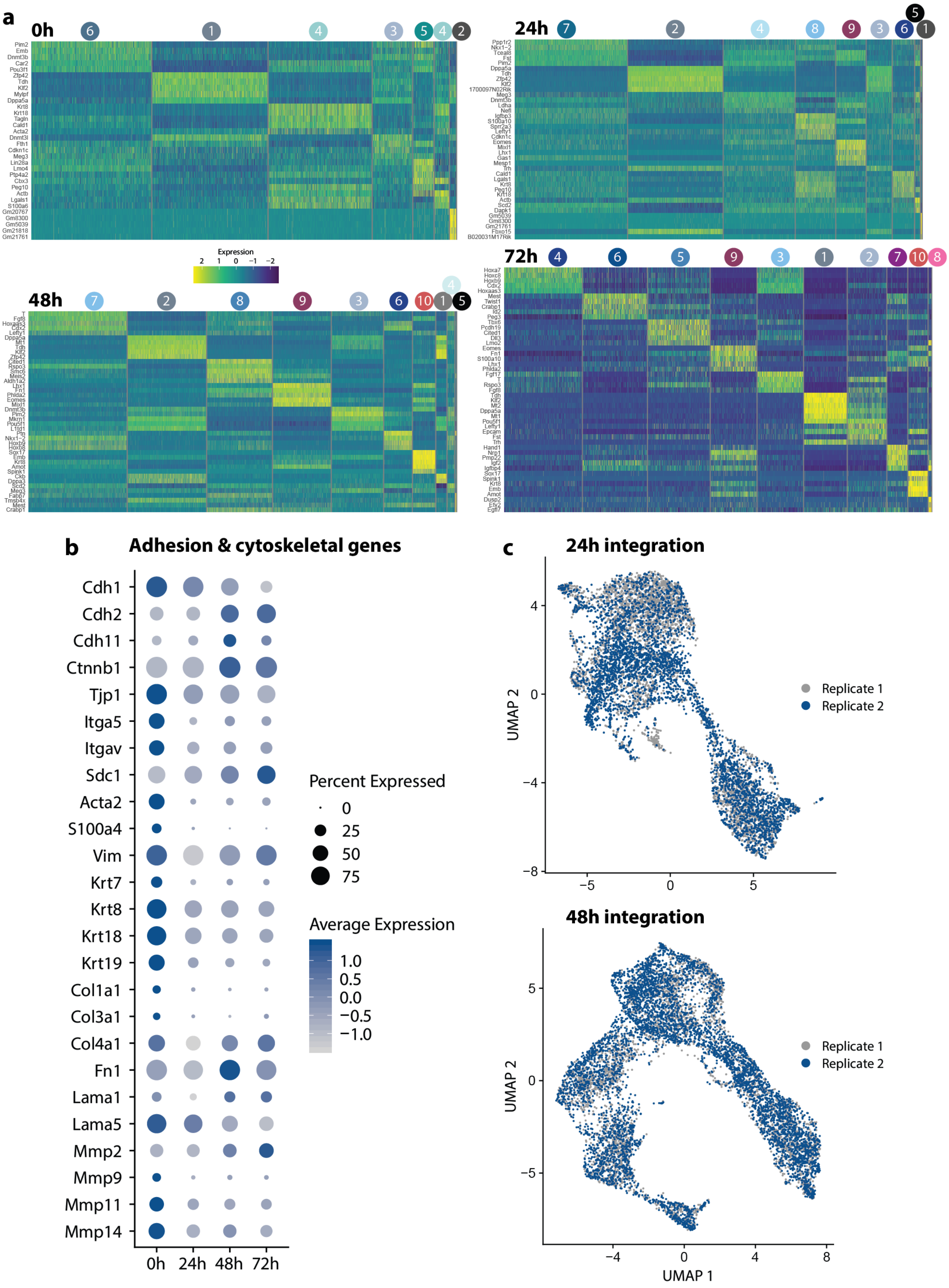


Supplementary Figure 2: **Single-cell transcriptomic analysis of early gastruloid development** *(**a**) Heatmaps showing the top 5 genes (if available) of UMAP clusters of mESC (0h) and gastruloid scRNA datasets from 24h to 72h AA. Cluster labels correspond to labels in Figure 2a. (**b**) Dot plots displaying expression of adhesion & cytoskeleton-related genes across each timepoint of the entire dataset. Size of circles denotes fraction of expressing cells and color indicates average scaled expression (Av. Expr.) levels. (**c**) UMAPs of gastruloid datasets at 24h and 48h AA color-coded by replicate demonstrating successful integration and reproducibility of cell types.*

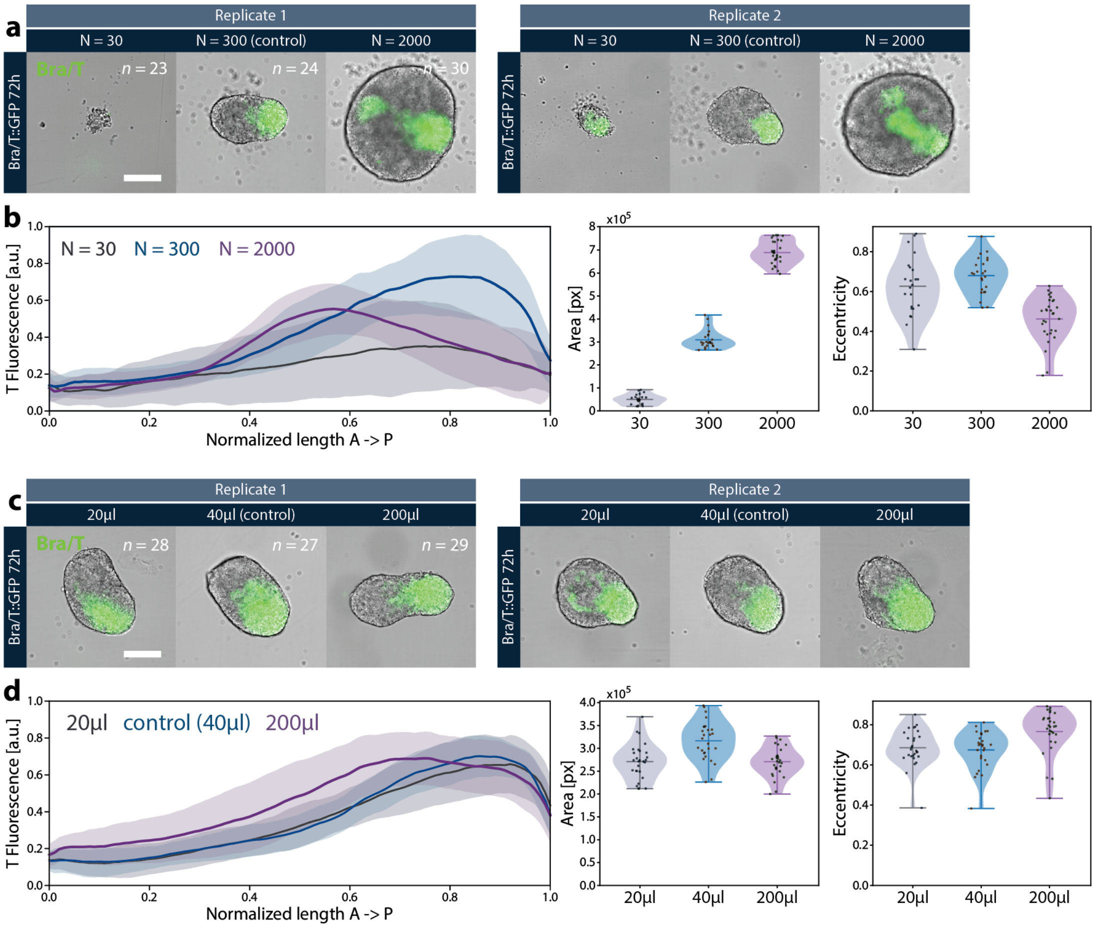


Supplementary Figure 3: **Exploring the regulative capacity of Bra/T polarization in gastruloids** *(**a**) and (**b**) Exemplary images and quantification of Bra/T fluorescence intensity along the AP axis as well as aggregate area and eccentricity for gastruloids grown from different starting cell (N) numbers demonstrating the size or cell number upper and lower limits for reproducible Bra/T polarization. (**c**) and (**d**) Representative images and quantification of Bra/T fluorescence intensity along the AP axis as well as aggregate area and eccentricity for gastruloids grown in different media volume to assess the importance of auto-secretion of signals for AP axis formation. For the 200µl condition, medium was additionally exchanged at 24h AA. Scale bars = 200µm.*

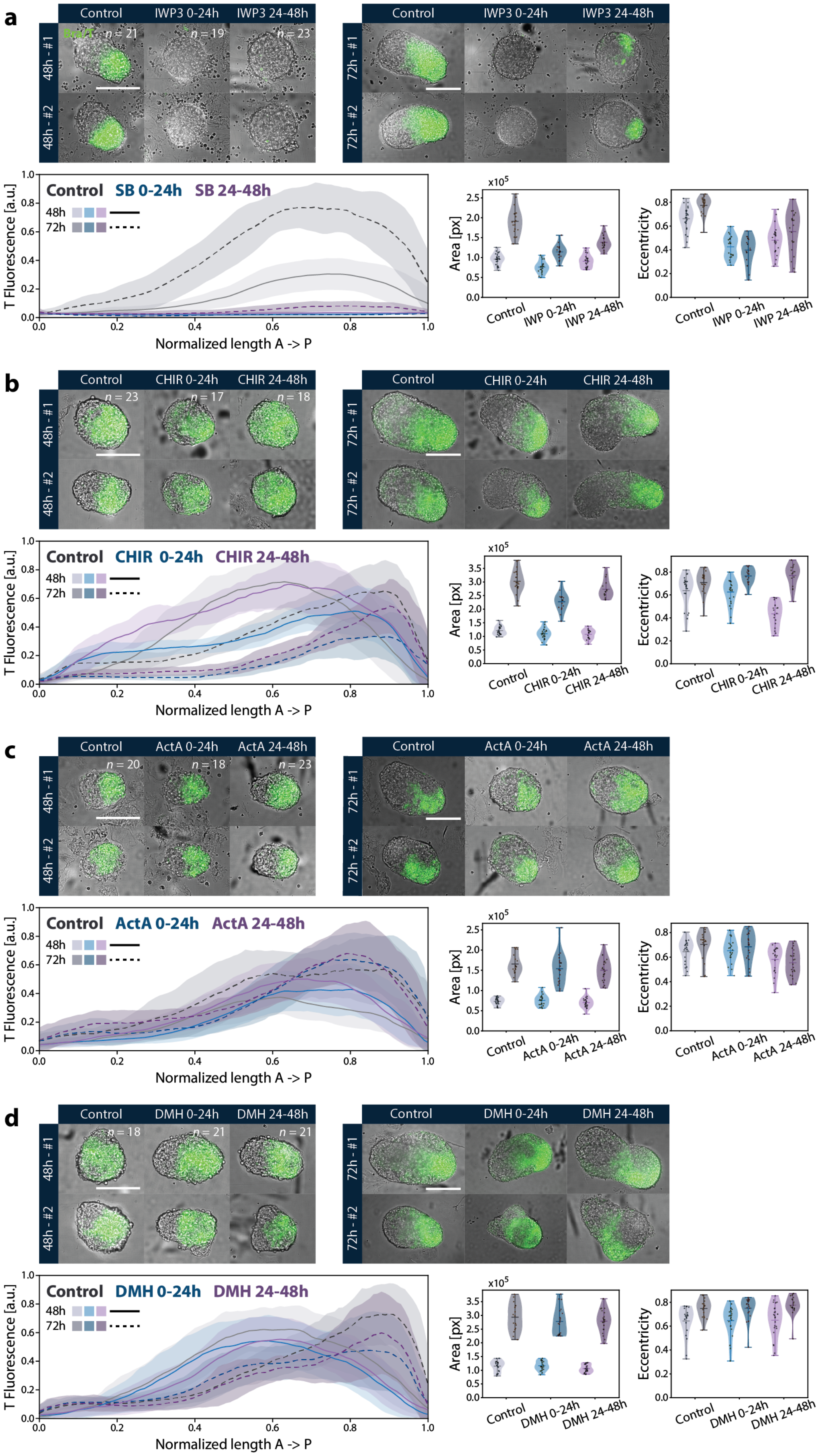


Supplementary Figure 4: *Caption on page 6.*

Supplementary Figure 4: **Assessing signalling requirements for Bra/T symmetry breaking**
*(**a**) - (**d**) Timed perturbations of key developmental signalling pathways Wnt via IWP3 (IWP) (**a**) and CHIR99 (CHIR) (**b**), Nodal/TGF-β via ActivinA (ActA) (**c**) as well as BMP via DMH1 (DMH) (**d**) to dissect their respective roles during Bra/T polarization in gastruloids. Representative images as well as dataset quantifications (Bra/T fluorescence intensity along the gastruloid's AP axis, area, eccentricity) are shown for each small molecule inhibitor corresponding to a pathway and for each inhibition time frame (0-24h AA and 24-48h AA, respectively). Scale bars = 200µm.*

Supplementary Video 1: **Onset of Bra/T expression in gastruloids**
*Timelapse imaging of Bra/T::GFP mESCs aggregated into gastruloids. Bra/T channel is displayed in green. n = 15.*

Supplementary Video 2: **Progression of Bra/T polarization**
*Light sheet live imaging of Bra/T::GFP gastruloids from 24h to 48h AA shows the Bra/T symmetry breaking or AP axis formation event. n = 3.*

Supplementary Video 3: **Bra/T expression dynamics upon MEK/ERK inhibition**
*GFP channel timelapse imaging from 48 to 72h AA of Bra/T::GFP gastruloids treated with MEK/ERK inhibitor PD03 from 24 to 48h AA depicts emergence of Bra/T expression as a band-like domain. n = 17.*

Supplementary Video 4: **Bra/T expression dynamics upon TGF-β/Nodal inhibition**
*GFP channel timelapse imaging from 48 to 72h AA of Bra/T::GFP gastruloids treated with TGF-β/Nodal inhibitor SB43 from 0 to 24 AA illustrates emergence of Bra/T expression slightly offset from the elongating tip. n = 14.*
